# Accurate staging of chick embryonic tissues via deep learning

**DOI:** 10.1101/2022.02.18.480991

**Authors:** Ian Groves, Jacob Holmshaw, David Furley, Matthew Towers, Benjamin D. Evans, Marysia Placzek, Alexander G. Fletcher

**Affiliations:** School of Mathematics and Statistics, University of Sheffield, Hicks Building, Hounsfield Road, Sheffield, S3 7RH, UK; School of Biosciences, University of Sheffield, Firth Court, Western Bank, Sheffield, S10 2TN, UK; Department of Informatics, School of Engineering and Informatics, University of Sussex, Falmer, Brighton, BN1 9RH, UK

**Keywords:** deep convolutional neural networks, data augmentation, hypothalamus, chick embryo, somites, wing bud

## Abstract

Recent work has indicated a need for increased temporal resolution for studies of the early chick brain. Over a 10-hour period, the developmental potential of progenitor cells in the HH10 brain changes, and concomitantly, the brain undergoes subtle changes in morphology. We asked if we could train a deep convolutional neural network to sub-stage HH10 brains from a small dataset (<200 images). By augmenting our images with a combination of biologically informed transformations and data-driven preprocessing steps, we successfully trained a classifier to sub-stage HH10 brains to 87.1% test accuracy. To determine whether our classifier could be generally applied, we re-trained it using images (<250) of randomised control and experimental chick wings, and obtained similarly high test accuracy (86.1%). Saliency analyses revealed that biologically relevant features are used for classification. Our strategy enables training of image classifiers for various applications in developmental biology with limited microscopy data.

**SUMMARY STATEMENT:** We train a deep convolutional network that can be generally applied to accurately classify chick embryos from images. Saliency analyses show that classification is based on biologically relevant features.

## INTRODUCTION

Developmental biology studies rely on the accurate staging of embryos, traditionally achieved with reference to simple morphological features described in conventional charts (Hamburger and Hamilton, 1951; O’Rahilly and Müller, 2010; Theiler, 2013). However, new approaches are enabling a greater temporal resolution of cellular and molecular events in developing embryos, and consequently researchers increasingly require more detailed staging systems (Newgreen and Erickson, 1986; Palmeirim *et al*., 1997; Boehm *et al*., 2011; Sáenz-Ponce *et al*., 2012; Musy *et al*., 2018).

Deep neural networks (DNNs) are increasingly used for image classification (LeCun *et al*., 2015) and are promising tools for staging embryos. Generally, DNNs require large training datasets for optimal performance (Deng *et al*., 2009; Thompson *et al*., 2020; Jacquemet, 2021). When trained on small datasets (100s-1000s of images), DNNs may exhibit poor performance on new data due to insufficient learning of general classifying features (Rosin and Fierens, 1995). Data augmentation techniques, involving image transformations, can improve generalisation by helping DNNs to reduce overfitting, increasing focus on class-defining image features and disregarding irrelevant features such as acquisition artefacts (Simard *et al*., 2003). In this way, DNNs have been used to classify embryonic developing systems from small datasets, where acquiring more images is impractical due to time, cost, or ethical considerations. For example, a DNN was used to stage zebrafish tailbuds as a model for posterior spinal cord growth (Pond *et al*., 2021). A second study trained a DNN to accurately classify zebrafish embryos as normal or malformed based on morphology, and demonstrated generalisation capability (Ishaq *et al*., 2017). However, these studies did not investigate how each DNN interpreted the datasets to successfully achieve classification. Saliency mapping, which highlights the image features used by a DNN classifier, shows promise in providing interpretability (Simonyan *et al*., 2014). By revealing the inner workings of high-accuracy DNN classifiers, saliency maps can help demystify their “black box” nature, facilitating their wider adoption in developmental biology.

In this study, we asked if we could train a DNN to successfully classify sub-stages within the HH10 chick brain. The embryonic chick benefits from a well-defined, precise, and detailed staging system that classifies embryos from HH1 to HH46 (Hamburger and Hamilton, 1951; Stern, 2018), but this classification can be insufficient for capturing temporal transitions that occur within individual stages. Our recent studies reveal rapid changes in gene expression in the ventral forebrain over HH10 that reflect the changing developmental potential of hypothalamic progenitor cells (Fu *et al*., 2017; Kim *et al*., 2022; Chinnaiya *et al*., 2023). Precise staging is therefore crucial to accurately study embryos during this period, particularly for targeted experiments of live embryos. While precise stages can be easily assigned after post-hoc post-fixation analyses such as *in situ* hybridisation (Kim *et al*., 2022; Chinnaiya *et al*., 2023) this is more challenging before fixation. More refined HH10 staging (HH10-, HH10, HH10+) traditionally relies on somite number (9, 10, 11 somites respectively), yet studies in Xenopus suggest that the head and body do not always develop synchronously (Sáenz-Ponce *et al*., 2012), and no study has asked, in the chick, whether somite number is an accurate predictor of brain development.

Here, we asked if we could train an DNN to accurately sub-stage HH10 chick brains from a small microscopy dataset of the heads of live chick embryos. A bespoke DNN performed optimally, achieving classification up to 87.1% test accuracy. Saliency analyses identified features of the brain typically used to classify the sub-stages. These included the same features used by experts to define the sub-stages, and an additional novel morphological feature. We then showed that the classifier could be re-trained to categorise growth-inhibited and normal wing buds, achieving a test accuracy of 86.1%. Here, saliency analysis revealed that the classifier used features that were not obvious to the human eye, demonstrating its applicability to uncharacterised specimens. Our accurate classifiers are valuable tools for chick embryo experimentation and our studies reveal how saliency analysis provides unbiased insights into predictive biological features.

## RESULTS

### Contemporary studies motivate an accurate sub-staging of the HH10 chick brain

In recent studies, in which we live-imaged chick embryos (Chinnaiya *et al*., 2023), we noted a continuous change in brain morphology during HH10 that could not be easily appreciated in fixed specimens with reference to conventional staging charts. The prosencephalon changes from a oval-shaped to a triangular-shaped structure as the optic vesicles widen, and the angle of the prosencephalic neck changes from obtuse to orthogonal and then acute (**Fig 1A-C, A’-C’**). Simultaneously, the hindbrain widens, the rhombomeres start to become distinct (**Fig 1A-C, A’-C’**), and a characteristic flexure forms in the prosencephalic ventral midline (indicative of the region where tuberal hypothalamic progenitor cells are generated) (Chinnaiya *et al*., 2023). Acutely-dissected HH10 embryos can be categorised into ‘early’ and ‘late’ based on these morphologies by experts with years of experience, but those with less experience can find this challenging (**Fig 1B-E, B’-E’**). We asked whether we could sub-stage early and late HH10 chick brains by counting somites. Unexpectedly, while head morphology did correlate with somite number at a population level, individual embryos with distinct brain morphologies could show the same number of somites (**Fig 1F**, **G**).

**Fig 1.**
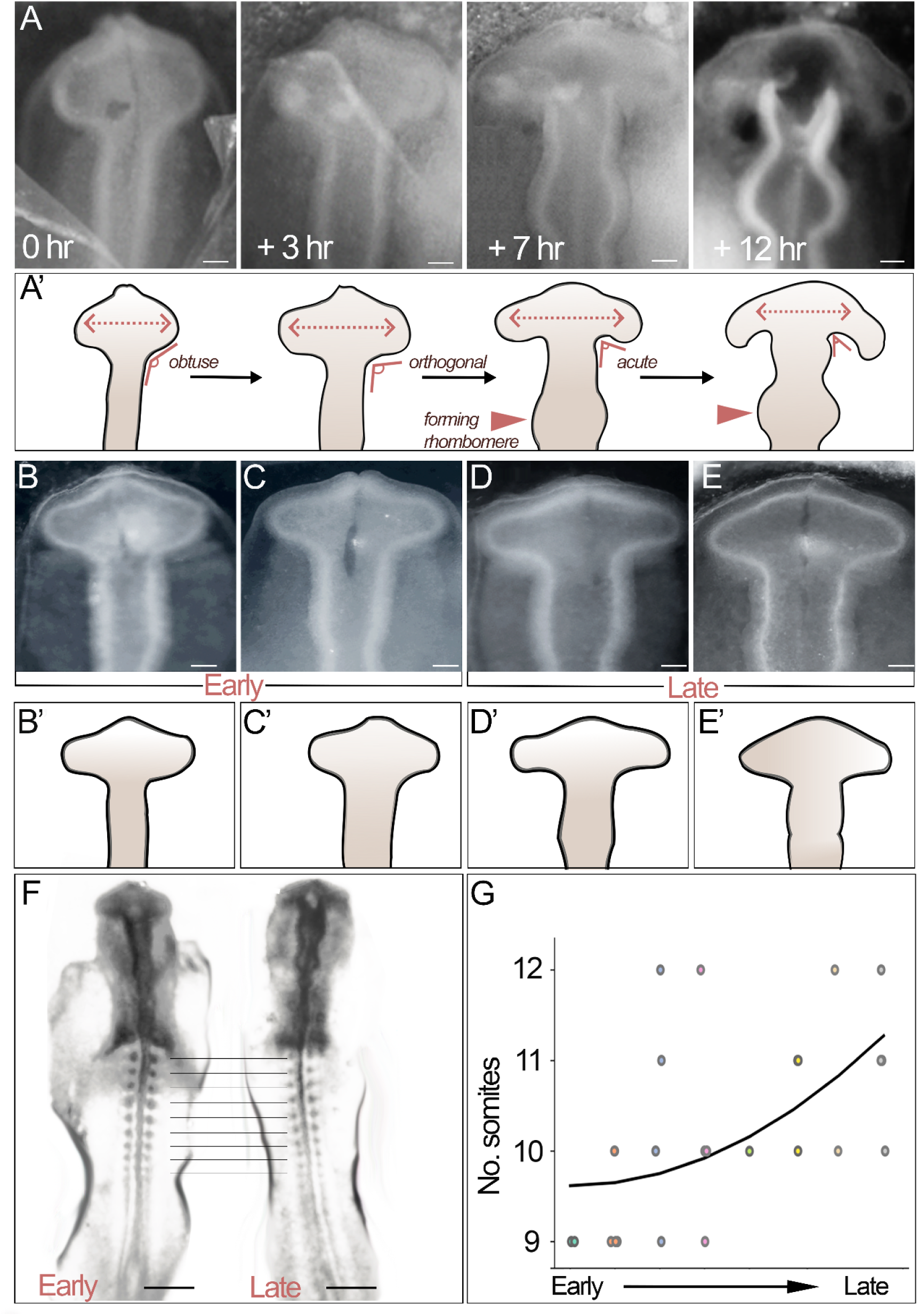
Somite number does not accurately predict brain development at HH10. (A) Live-imaging reveals the rapid morphological changes in the brain as a HH10 embryo (first 3 panels) develops to HH11 (4th panel) (n=6). (A’) Schematics of images shown in (A) pointing to key morphological features used for classification: prosencephalic width; angle of prosencephalic neck; shape of developing hindbrain. (B-E’) Brightfield views of individual embryos, with early prosencephalic morphology (B, C and schematics B’, C’) or late prosencephalic morphology (D, E and schematics D’, E’). (F) Brightfield views of embryos with distinctive brain morphologies (early and late), but similar somite numbers. (G) Number of somites in HH10 embryos (n=22), ordered according to head morphology from early to late. Scale bars: (A-E) 100μm; (F) 500μm.

The accurate categorisation of the HH10 prosencephalon into early versus late substages is important, because over this period, cells – at least those in the ventral prosencephalon – rapidly change in character and developmental potential. In HH10 embryos with an ‘early’ prosencephalic morphology, *SHH* is co-expressed with *BMP7*, marking hypothalamic floor plate-like (HypFP) cells (**Fig 2A, A’**), but in embryos with a ‘late’ prosencephalic morphology, *SHH* extends more anteriorly than *BMP7*, marking progenitors that will go on to generate tuberal hypothalamic neurons (**Fig 2B, B’**) (Kim *et al*., 2022; Chinnaiya *et al*., 2023).

**Figure 2.**
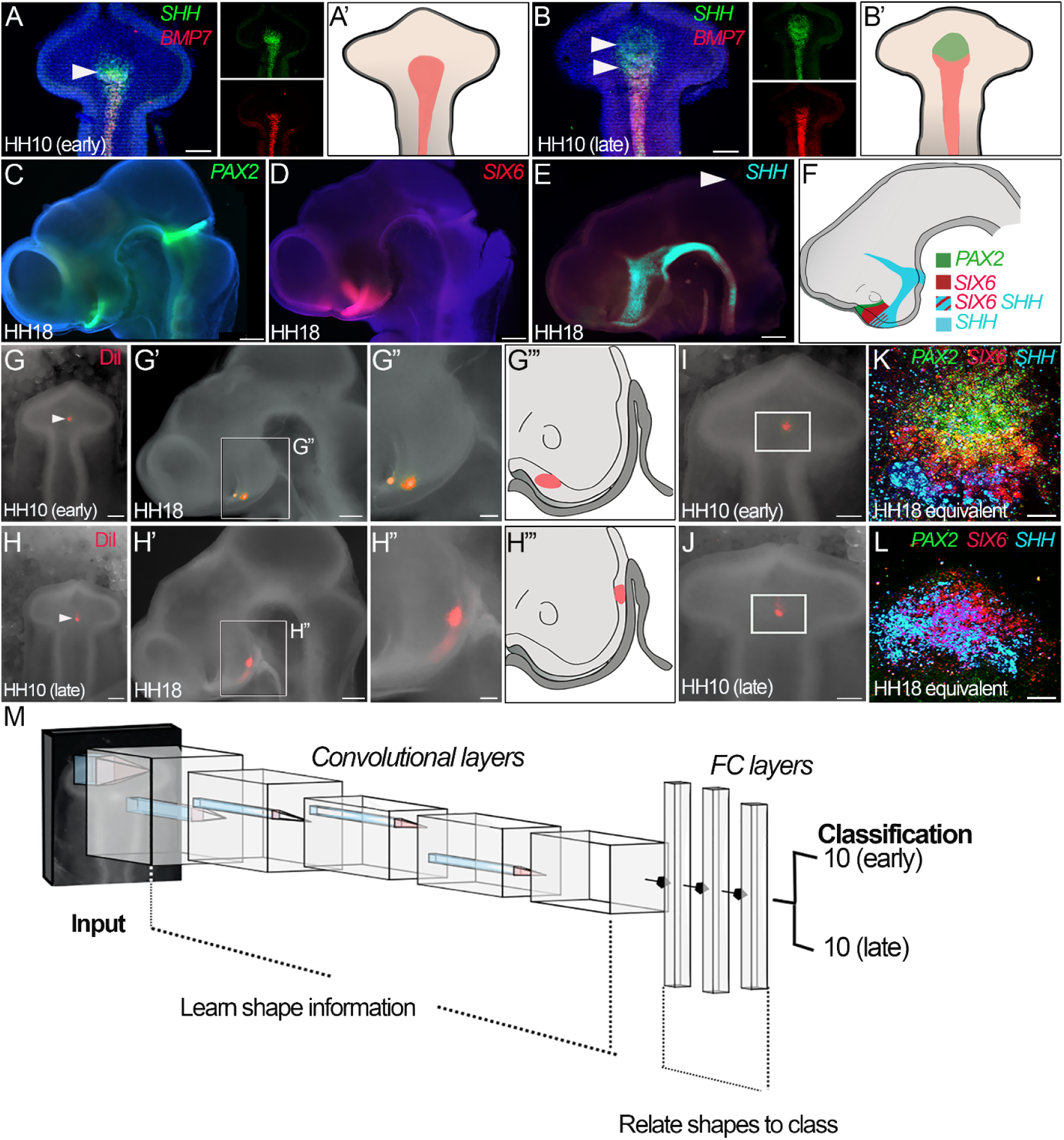
Changing developmental potential in the HH10 chick brain. (A-B’) Ventral wholemount views of isolated brains from HH10 embryos with ‘early’ (A) and ‘late’ (B) morphologies after HCR to detect expression of *SHH* and *BMP7.* Anterior-most HypFP cells co-express *SHH* and *BMP7* and extend to a characteristic flexure in the ventral midline (white arrowheads in A, B). Tuberal progenitors express *SHH* and are readily detected in embryos with a ‘late’ prosencephalic morphology (green arrowhead in B) (n>20). Expression patterns summarised schematically in (A’, B’). (C-F) Side-views of HH18 heads after HCR to detect expression of *PAX2* (C), *SIX6* (D) or *SHH* (E), which mark cells with different positions along the anterior-posterior axis (n>10), summarised schematically in (F): optic stalk (*PAX2)*; anterior tuberal neurogenic progenitors (*SIX6);* anterior tuberal and supramammillary progenitors, ZLI and floor plate (*SHH);*. (G-H”’) Targeted injection of DiI into nascent tuberal progenitors in HH10 embryos with ‘early’ (G) or ‘late’ (H) prosencephalic morphologies (n>10 each). Cells targeted in an ‘early’ embryo fate-map to the anterior-most tuberal region, just posterior/ventral to the optic stalk (G’, G”, shown schematically in G”’); cells targeted in a ‘late’ embryo fate-map to more posterior tuberal regions, overlying Rathke’s pouch (H’, H”, shown schematically in H”’). (I, J) Same embryos as in (G, H), depicting regions explanted (boxed), encompassing anterior-most HypFP cells and adjacent regions. (K, L) Explants taken from ‘young’ or ‘old’ HH10 embryos, cultured for 48 hours, and analysed by HCR to detect *PAX2* (green), *SIX6* (red), and *SHH* (cyan) (n=5 each). (M) Schematic indicating the operation of a convolutional neural network for sub-stage classification. FC: Fully connected. Scale bars: (A, B) 100μm; (C-E) 250 μm; (G, H) 100μm; (G’, H’) 250μm; (G’’, H’’) 100μum; (I-L) 100μm.

Importantly, the changing gene expression profile at HH10 reflects changing developmental potential. Fate-mapping studies of the ventral prosencephalon, from HH10 to HH18 when distinct progenitor subsets can be identified based on position and molecular profile (**Fig 2C-F**) (Fu *et al*., 2017; Chinnaiya *et al*., 2023), have shown that tuberal progenitors are sequentially generated from HypFP cells, with those born earliest lying close to the optic stalk and those born later lying above Rathke’s pouch. Thus, HypFP cells targeted in chicks with ‘early’ versus ‘late’ prosencephalic morphology fate-map to sequentially more posterior parts of the tuberal hypothalamus at HH18 (**Fig 2G-G’**, **H-H’**). Finally, prosencephalic morphology is an accurate predictor of cell specification. When prosencephalic tissue of equivalent size and region (using the prosencephalic neck as a reference point) is explanted from HH10 embryos (**Fig 2I, J**) and cultured to a HH18 equivalent, explants taken from an ‘early’ prosencephalon express optic stalk (*PAX2*) and tuberal progenitor markers (*SIX6* and *SHH)* (**Fig 2K**), whereas explants taken from a ‘late’ prosencephalon express only tuberal progenitor markers (*SHH* and *SIX6*) (**Fig 2L**).

Together, these studies demonstrate the importance of accurately staging the HH10 brain.We therefore asked if we could accurately stage live HH10 embryonic brains using an automated classification tool.

### Fine-tuning the ResNet50 architecture classifies sub-stages of HH10 with up to 75% accuracy

In order to train a classifier, we used our expertise to group images of HH10 embryos into the two sub-stages (early and late). We first asked if unsupervised machine learning methods could be used to classify these images. We tested clustering approaches including principal component analysis and *k*-means (Ding and He, 2004) using both raw images and features extracted using conventional Haralick ‘texture’ (Haralick *et al*., 1973). We then tested traditional supervised classifiers (Amancio *et al*., 2014), in particular, random forest classifier (RFC), support vector machine (SVM), and *k-*nearest neighbours (KNN). We were not able to train a sufficiently accurate classifier through any of these approaches, achieving the highest individual and highest average validation accuracies of only 54.8% (RFC) and 38.3% (KNN), respectively, through supervised classifiers (Table S1, Figs S1-S2).

Therefore we developed a strategy for training a deep convolutional neural network (DCNN) based classifier. DCNN classifiers have proven particularly powerful in image classification, as they contain convolutional layers which learn filters representing important shape information contained in the images (LeCun *et al*., 2015). First, we determined suitable data preprocessing approaches (see *Materials and Methods*). Next, we augmented the dataset through image transformations (**Fig S3**), which expanded the number of datapoints for training, and normalised skewed image features such as subject orientation that are unlikely to be important for classification. We examined the benefits of various image augmentations, setting rotations as our baseline augmentation (Ishaq *et al*., 2017). Finally, we implemented a cross-validation approach to improve our DCNN’s generalisation by systematically changing the data in the training and validation sets. In cross-validation, we varied both the training data (used for fitting the DCNN) and the validation data (used to evaluate the generalisation performance of the learned features). However, we fixed the test dataset to allow fair comparisons when ultimately evaluating the classifier for an unbiased estimate of generalisation (see *Materials and Methods*).

Using these approaches, we evaluated the viability of transfer learning to train a DCNN classifier, a commonly used approach for dealing with small datasets (Kora *et al*., 2022). Specifically, we asked if we could use the pre-trained DCNN classifiers, InceptionV3 (Szegedy *et al*., 2015) and ResNet50 (He *et al*., 2016), both of which have achieved high classification accuracies on a database comprising over 14M general images.

We re-trained these models on our brain dataset. Generally, InceptionV3 performed poorly, with average accuracies in the range of 47-52% across the various augmentation regimes (**Table S2**). ResNet50 performed better (average accuracies in the range of 50-70%), but the highest individual model accuracy achieved was still only 75.9% (baseline and Gaussian blur regime). Additionally, this regime achieved the second lowest standard deviation (6.8%), an important metric in light of a limited dataset (**Table S2**). Taken together, these results suggest that training with commonly-used image classification architectures is not effective in classifying small microscopy datasets.

### A bespoke neural network classifies brain sub-stages with up to 87% accuracy

We therefore next asked whether we could improve classification accuracy beyond that achieved by Inception/ResNet50 by designing a bespoke DCNN. Our investigations using Inception V3 and ResNet50 had revealed that performance could be substantially improved by data preprocessing, and the selection of particular augmentation regimes. In addition, classifier performance can be improved by optimising parameters that are set prior to training. These hyperparameters comprise the overall computational architecture of the network, including the number of computational units, and the rate at which these units update their connection weights – the learning rate. Hyperparameters are typically optimised via systematic (LeCun *et al*., 1998) or random (Bergstra and Bengio, 2012) search. Bayesian optimisation techniques are increasingly used, with a probability model informing which values to test (Shin *et al*., 2020). An open question then, when training DNNs on microscopy images, is how best to exploit the combination of hyperparameters (e.g. network architecture) and data augmentation techniques to suit typically small datasets in developmental biology.

We chose to construct a model with a wide, VGG-16 block-style architecture (**Fig 2M**), which has been successful in image classification (Simonyan and Zisserman, 2014). We used Bayesian optimisation and empirical selection to tune the hyperparameters (**Table S4**). We then determined the most useful and robust augmentation regimes (**Table 1**, brain dataset). Overall our bespoke DCNN with our baseline augmentation regime performed well, surpassing our best ResNet50 results (average test accuracy of 73.5%). Better still, across the training process, each augmentation resulted in a higher validation accuracy than the baseline augmentation alone (rotation, see above), our best performing augmentation set being ‘baseline & shear’ (83.9% test accuracy). We also tested the efficacy of Möbius transformations, a class of geometric mappings that have proven successful in other limited data contexts (Zhou *et al*., 2021) but are untested for microscopy image classification. We reasoned that Möbius transformations could introduce the DCNN to common microscopy artefacts, e.g. tissue bending during sample preparation. However, our baseline and Möbius transformations performed more poorly than the baseline alone (**Table S3**, average accuracy: 66.1%). Augmenting only 10% of the data with Möbius transformations also decreased validation accuracy below our baseline results (**Table S3**, average accuracy: 75.9%).

**Table 1.**
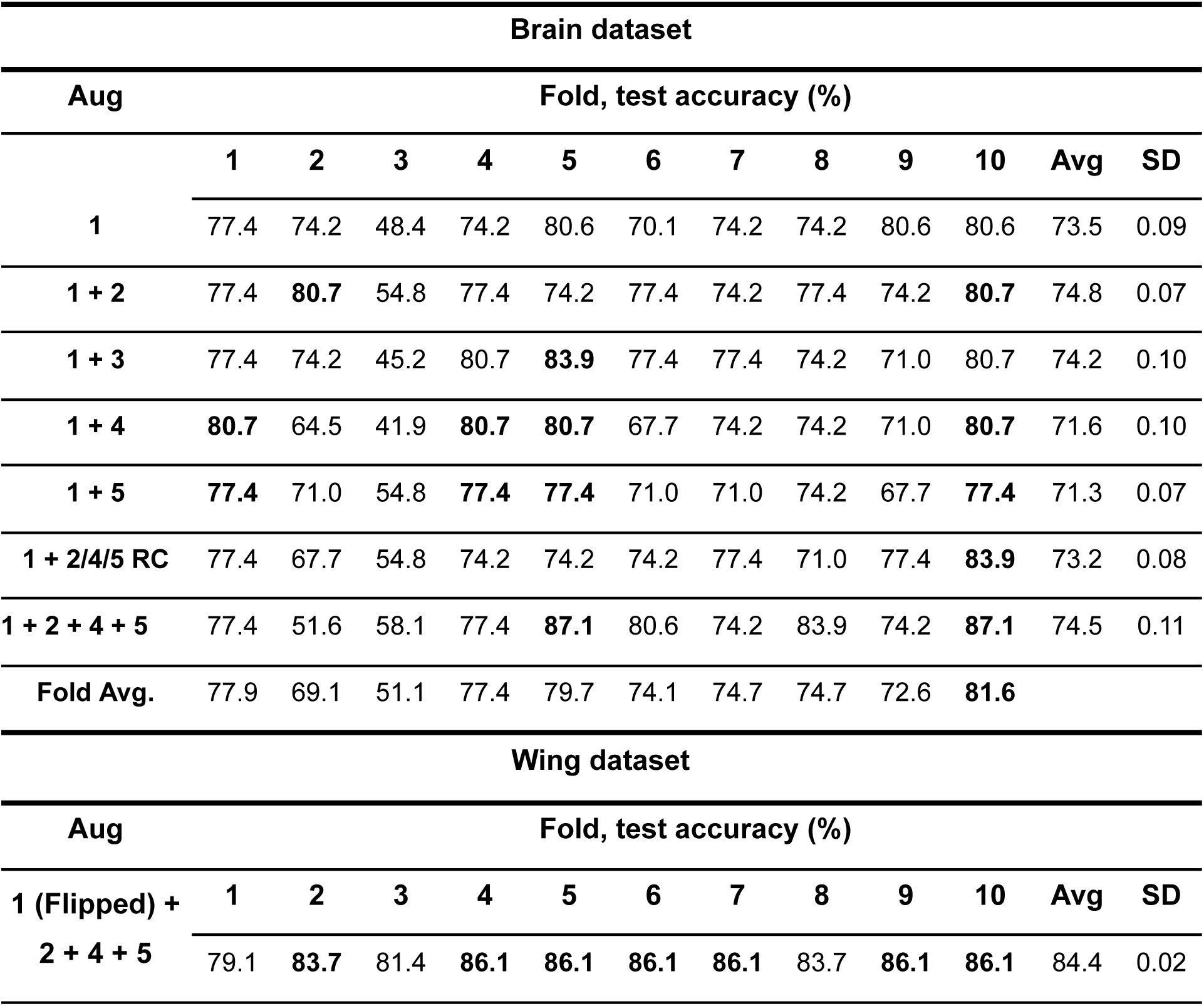
Augmentation exploration of the brain and wing datasets. For each dataset, we used nested cross validation, utilising *k*-fold cross validation for model tuning/augmentation selection and evaluating on an independent held-out dataset. The individual fold test accuracies achieved by each network are shown in columns 1-10, followed by their averages (Avg) and standard deviations (SD). As a baseline processing step, all images were rotated 15 times, at equally spaced degrees. For the brain dataset, we tested different augmentations on top of this baseline. For the wing dataset, we tested the combined augmentation regime that gave the highest test accuracy for the brain dataset. Augmentations (Aug): (1) baseline – rotation and (wing only) flip; (2) shear; (3) crop; (4) Gaussian blur; (5) cutout; (1+ 2/4/5 RC) random combination of baseline, Gaussian blur, and cutout.

Having confirmed that additive pairwise augmentations are useful (e.g. baseline rotation & Gaussian blur; baseline rotation & shear), we next asked whether more sophisticated combinations would further test accuracy: a (random) choice regime and a combined regime (**Table 1**). The first of these was pairwise as before, but the augmentation on top of baseline was chosen at random, so that each image had two augmentations applied. In the second, the combined regime, every transformation was applied to each image. In both cases, training a model using these combinations improved performance. We identified an informed combined regime which resulted in substantially higher test accuracies (**Table 1**, brain dataset model 10: 87.1%), i.e. in which the network had learned ‘difficult’ features of the images. We suggest that this regime is optimal when dealing with small datasets that exhibit high variability in DCNN training.

In summary, we constructed a bespoke DCNN that was substantially better suited to classifying HH10 brain substages than ResNet50 or InceptionV3.

### The re-trained DNN classifies chick wings with up to 86% accuracy

We next investigated whether the convolutional layers from our highest-scoring model (**Table 1** model 10; 87.1% test accuracy), and our preprocessing and data augmentation approach could be applied to a second, similarly sized microscopy dataset, using previously published data comprising 269 images of HH24-HH28 chick wings (Towers *et al*., 2008). In contrast to the HH10 brains, the wings are rather amorphous, and are not easily classifiable by the HH staging system.

The wing dataset comprised images from embryos in which a control bead, or a Trichostatin A-bead had been implanted at HH20, the embryos developed for up to 56 hrs, and then analysed for expression of *SHH*. Images were divided into two categories: ‘control’ (representing normal wing development) and ‘treated’. Trichostatin A transiently inhibits growth and leads to morphological changes in the wing bud, making it difficult to assign a HH stage. We asked whether the DNN could categorise wing buds on the basis of the drug-induced morphological changes, regardless of their presumptive developmental stage.

The training regime was similar to that used for brain classification, but with one additional augmentation: we included images that were flipped along the horizontal axis. This was motivated by the experimental design, where right wing buds were treated and left wing buds were left as control (Towers *et al*., 2008). Introducing flipped images was essential because the classes were always oriented in one direction, so DCNNs that were trained without the flipped versions would overfit substantially, with highly exaggerated accuracy results. To ask whether the shape features (‘filters’) learned by the DCNN during brain dataset training could be useful for other morphological problems we froze the feature extractors (filters) learnt in the convolutional layers learned on the brain dataset (Zeiler and Fergus, 2014) and trained only the fully connected layers at the end of the network (**Fig 2M**, FC): the latter learn the relationship between the extracted shapes and the classification (Yamashita *et al*., 2018).

We found that the test accuracies achieved were generally even higher than for the brain classification (average test accuracy 84.4%; highest accuracy on any individual model 86.1%: **Table 1**, wing dataset). Thus our brain dataset based DCNN, trained via a strategy of reasoned data augmentations, extended well, classifying another limited microscopy dataset of developing wings with high accuracy with minimal modifications to the training pipeline.

### Saliency maps identify biologically relevant class-specific features

Having trained two accurate DCNN classifiers on two separate datasets, we next performed saliency analysis on each classifier to determine the image region(s) to which it was sensitive. In the case of the brain dataset, we asked if the DCNN had recognised the relevant features used by experimentalists. For the brain dataset, we selected the best-performing classifier from **Table 1** (brain dataset, model 10) and generated saliency maps for test images across each sub-stage (**Fig 3A-E**), as well as maps in which we had filtered out low-level activations (**Fig 3A’-E’**), and scored each image in the test dataset based on the areas with high levels of attention.

**Fig 3.**
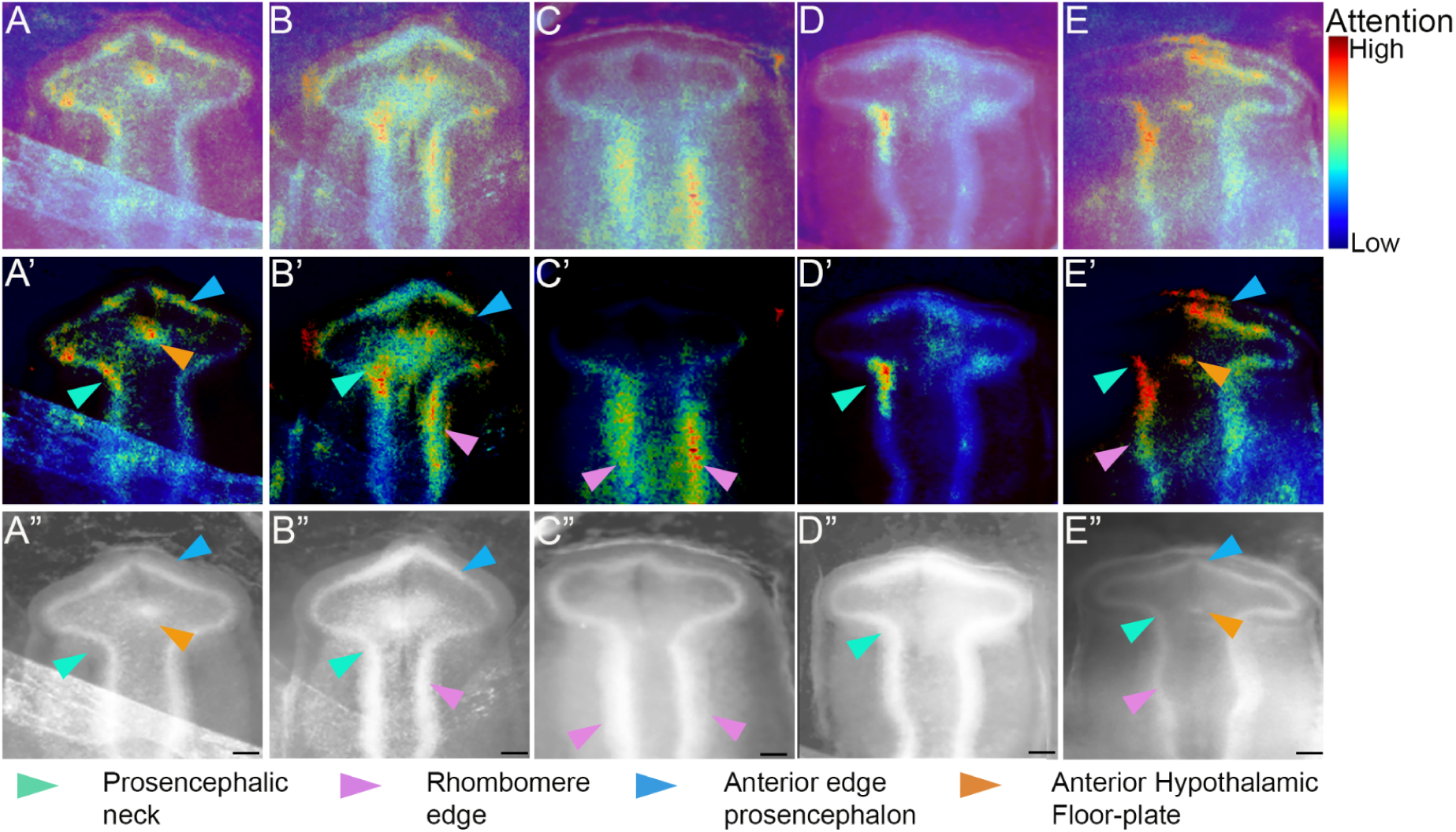
Saliency maps of HH10 (early) and HH10 (late) sub-stages highlight defining morphological features. (A-E) Saliency maps of HH10 (early) embryos (A-C), and HH10 (late) embryos (D,E) generated by the highest performing (87.1% test accuracy) bespoke classifier (**Table 1**, brain dataset, model 10). None of the images was used in training/validation of the DCNN. (A’-E’) As in A-E but with low level saliency pixels filtered out. (A’’-E’’) corresponding test input images made grayscale and with brightness/contrast normalised. Coloured arrowheads point to regions of high attention. The entire test dataset is scored with key morphological regions counted. Prosencephalic neck: 71%; Rhombomere edge 71%; Anterior edge of the prosencephalon: 50%; anterior hypothalamic floor plate: 33%. Note that the same regions of pixels can be relevant to both classes in a DCNN. For example, if the shape of the rhombomeres is crucial for distinguishing between the 10 (early) class and 10 (late) sub-stages, then the network could focus on the rhombomeres in saliency maps for both classes. Scale bars: 100μm.

For both brain substages, the most salient regions were those used by the human experts to initially classify the data: the prosencephalic neck and the rhombomere edges (**Fig 3A** cyan and magenta arrowheads); 71% of the test dataset had high activation in these regions. Additionally, 33% showed focus on the characteristic flexure in the prosencephalic ventral midline where the nascent tuberal hypothalamus is located (**Fig 3**, orange arrowheads). Additionally, 50% of the maps showed focus on the anterior edge of the prosencephalon (**Fig 3**, blue arrowheads), a feature not accounted for in the initial classification, but which could potentially reflect the changing angle of the prosencephalic neck. Taken together, these results show that the model has learned new as well as previously characterised biologically relevant class-defining features.

We next generated saliency maps from the wing classifier (**Fig 4A-H**), asking whether classification was by attention to obvious features, such as *SHH* expression/limb size, or by another means. As with the brain saliency analysis, we again filtered high levels of activation (**Fig 4A’-H’**) and scored the saliency maps with reference to morphological landmarks (**Fig 4A’-H’,** arrowheads). The saliency maps did not focus attention on any individual feature. The most consistent regions of high activation were the anterior margin of the wing, where 69% of the maps showed a focus (**Fig 4B’, C’, E’, G’, H’**, magenta arrowheads), and the distal edge of the wing, where 50% of the maps showed a focus (**Fig 4B’, C’, E’, H’**, blue arrowheads). There were three more morphological features with obvious activation: the posterior margin (the focus of attention in 33% of the saliency maps (**Fig 4C’, G’,** red arrowheads), the ‘shoulder’ region where the anterior edge of the wing meets the trunk (24%) (**Fig 4B’, D’, F’, G’**, green arrowheads), and the proximal edge, spanning the anterior-posterior axis (12%) (**Fig 4B’, F’**, wing width anterior-posterior). Surprisingly, the classifier did not consistently pay attention to the presence of *SHH* expression; (**Fig 4F’**’, orange arrowhead: only 17% images showing such focus). Overall, the saliency analyses reveal how classification can been made through learning of features by the DCNN that are not immediately obvious to the experimentalist.

**Fig 4.**
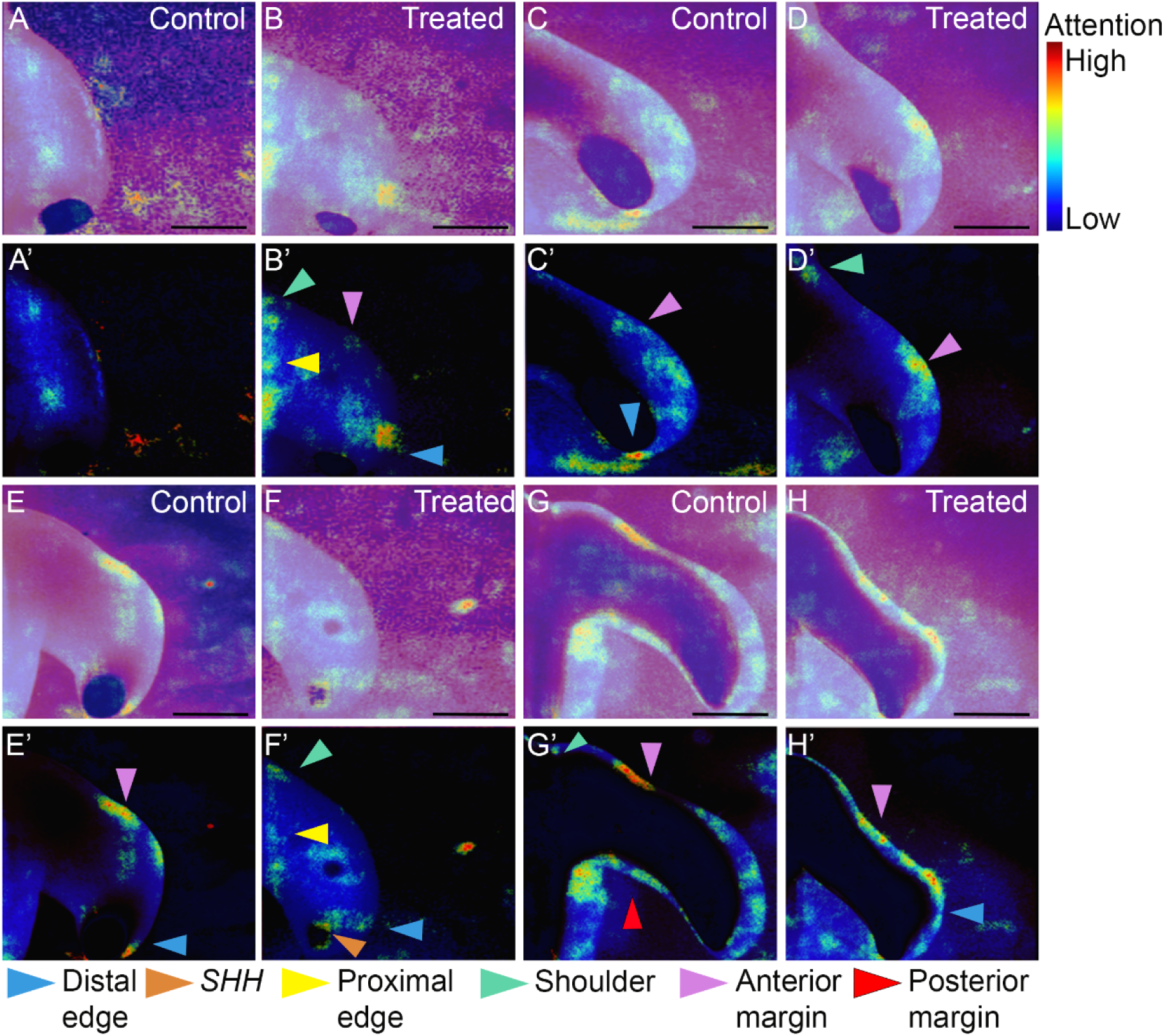
Saliency maps identify important morphological features in the classification of developing chick wings. (A-H) Saliency maps of control (A, C, E, G) and TSA growth inhibited (B, D, F, H) wings paired according to approximate wing size generated by the 86% test accuracy bespoke model (**Table 1**, wing dataset, model 4) on an independent (not using DCNN training/validation) test dataset. (A’-H’) As in A-H but with the low level saliency pixels filtered out. Input images were converted to grayscale with histogram normalisation applied. The saliency maps in the entire test dataset are scored according to morphological features: shoulder (green arrowheads), proximal and distal edges of the wing (yellow and /blue arrowheads respectively), and anterior and posterior wing margins (magenta and red arrowheads respectively). Scale bars 500μm.

In general, DCNNs are not directly interpretable and are often considered ‘black box’ solutions, and it remains an open challenge when training DCNNs as to how best to interpret their output. Taken together, these results show how saliency analysis helps our interpretation of how the classifiers make decisions, including highlighting unforeseen morphological changes. Our findings illustrate how rigorous examination of classifier attention can give insight into data processing and augmentation efficacy and into the features of the embryo which most determine the classification.

## DISCUSSION

Classifying embryos into discrete stages is challenging due to the continuous nature of development. Advances in high-resolution methods, including imaging and scRNA-seq (Cutrale *et al*., 2019) suggest the need to stage embryos more accurately than is possible through traditional staging guides. Here, we demonstrated the biological imperative of sub-classifying the HH10 brain, and then asked whether this could be achieved through machine learning methods. Neither unsupervised nor traditional supervised computational methods were able to accurately classify brains in a manner that reflected their developmental stage. By contrast, our trained DCNN classifier was able to accurately classify the HH10 brain, through a focus on subtle morphological changes that reflect and extend beyond human expertise.

A consensus in the field of deep learning is that for small datasets, transfer learning using open-source models (e.g. Inception V3 and ResNet50, both trained on huge general datasets) can be used effectively. For instance, ResNet50 has been used successfully in other biomedical fields (Baltruschat *et al*., 2019). We found in our case that training using InceptionV3 was not effective. By contrast, ResNet50 performed surprisingly well (up to 75.9% accuracy) when re-trained on our data. However, this performance leaves room for improvement so we asked whether a bespoke network trained from-scratch (randomly initialised, rather than pretrained) would achieve even higher classification accuracies.

For training a model from-scratch on a small dataset, a main consideration is avoiding overfitting. Previous deep learning efforts to classify microscopy images in developmental biology have focused on hyperparameter optimisation (Pond *et al*., 2021) and rotational augmentations (Ishaq *et al*., 2017). By contrast, here we performed a thorough and systematic exploration of a wide variety of data processing and augmentation regimes. Importantly, our data augmentations help remove biases towards irrelevant features such as illumination, orientation, focus, size, and colour, which may vary between images. Moreover, these augmentations could be assisting the network to focus on (a) true sub-stage characteristic(s), rather than features arising from biological inter-sample variation. We found that model performance depended strongly on data augmentation, with a combination of individually successful augmentations proving most effective. Overall, our bespoke network achieved high classification accuracy of the chick brain (87.1%). To extend the application of our classifier, we applied similar augmentation regimes and fine-tuned our brain classifier on a wing dataset, achieving similarly high classification accuracies.

We used saliency maps to identify the image regions to which our classifier was sensitive. In the case of the brain dataset, the classifier was most sensitive to the prosencephalic neck, the rhombomeres and the ventral prosencephalic flexure – precisely those regions used by the human experts to initially classify the data. This indicates that classification by the DNN is made on the basis of biologically relevant features, boosting confidence in the efficacy of our augmentations, and the classifiers’ performance generally. The brain classifier identified one further region – the anterior prosencephalic edge – that had not featured in the initial classification. Retrospective analyses of the images reveal that, indeed, the slope of the anterior prosencephalon alters through HH10, as the optic vesicles lengthen. Therefore, the DCNN can provide insight into novel classifying features.

In the case of the wing classifier, we initially hypothesised that the DCNN would primarily rely on ‘simple’ features like overall size or the expression profile of *SHH* to categorise control and treated wings. Surprisingly, saliency analysis revealed that the DCNN paid little attention to these metrics and instead focused on other morphological characteristics. The simplicity of the wing bud’s structure, combined with the classifier’s emphasis on specific edges and regions, suggests that the images contain important information regarding subtle morphological differences between control and treated embryos. This illustrates how post-hoc classifier analysis can motivate new biological hypotheses. For example, the drug Trichostatin A is thought to inhibit growth through cell cycle arrest and apoptosis (Bouyahya *et al*., 2022), which suggests that these processes may be integral to correctly shaping the limb, warranting further investigation.

Overall, our results illustrate the utility of saliency analysis in interpreting image classifiers for developmental biology, similar to other biomedical fields (Baltruschat *et al*., 2019; Panwar *et al*., 2020). The use of saliency methods will encourage confidence in non-specialists to use DCNN-based classifiers. Further work could expand the dimensions of the images used. We used 2-D morphological profiles to train our classifiers. Extending this with 3-D fluorescent images, which are increasingly used in developmental biology, could provide a richer amount of information to the model and result in a more robust, accurate classifier. Additionally, including gene expression data into the actual training process for the brain classification could couple our sub-stages to biological mechanism(s). Importantly, our freely available pipeline extends naturally to developmental datasets with different problems or classes. Our chosen strategy will therefore allow image classifiers to be trained for other biological systems with limited microscopy data. Our DCNN provides a tool to stage embryos at greater temporal resolution than conventional staging systems, offers the potential to compare embryos of different species, and to assist experienced researchers studying unconventional or emerging experimental organisms in developing staging systems for these organisms.

## MATERIALS AND METHODS

### Chicks

Fertilised Bovan Brown eggs (Henry Stewart & Co., Norfolk, UK) were used for all studies, which were performed according to relevant regulatory standards (University of Sheffield).

### Neural tube isolation, explant dissection and culture

HH10 neural tubes were isolated from surrounding tissue by dispase treatment, as previously described (Ohyama *et al*., 2005). Explants were isolated from HH10 embryos after dispase treatment and cultured, as previously described (Ohyama *et al*., 2005), then processed for *in situ* hybridization chain reaction (HCR) as below.

### Hybridization chain reaction

Embryos, neural tubes or explants were fixed in 4% paraformaldehyde, dehydrated in a methanol series and stored at –20°C. Hybridization chain reaction (HCR) v3.0 was performed using reagents and protocol from Molecular Instruments Inc. Samples were preincubated with a hybridization buffer for 30 min and the probe pairs were added and incubated at 37°C overnight. The next day samples were washed 4 times in the probe wash buffer, twice in the 5xSSC buffer, and preincubated in Amplification buffer for 5 min. Even and odd hairpins for each gene were snap-cooled by heating at 95°C for 90 sec and cooling to RT for 30 min. The hairpins were added to the samples in Amplification Buffer and incubated overnight at RT in the dark. Samples were then washed in 5xSSC and counterstained with DAPI.

### Fate mapping

Fate-mapping studies were a retrospective analysis of previous work (Fu *et al*., 2017; Chinnaiya *et al*., 2023).

### Fluorescent image acquisition

Fluorescent images were taken on a Zeiss Apotome 2 microscope with Axiovision software (Zeiss) or Leica MZ16F microscope or Olympus BX60 with Spot RT software v3.2 or Nikon W1 Spinning Disk Confocal with Nikon software. Images were processed and digitally aligned using Image-J (FIJI) and Adobe Photoshop 2021.

### Data acquisition

Ground-truth data used for training and validating classifiers comprised bright-field and phase-contrast microscopy images of HH10 (9-12 somite) chick embryos (brain data) and HH18-24 (wing data), including published and unpublished data. Images were acquired using an Olympus BX60 microscope, a Zeiss AxioImager.Z1 microscope, and a Leica dissecting microscope at 4x or 10x magnification. The brain dataset comprised 152 images (70 ‘early’ and 82 ‘late’) and was acquired as outlined in (Fu *et al*., 2017; Chinnaiya *et al*., 2023). Brain training data were labelled into two sub-stages, ‘early’ and ‘late’ assigned according to the overall shape of the prosencephalon, the angle of the posterior prosencephalon relative to the prosencephalic neck, the optic vesicle and rhombencephalon shape. The wing dataset contained 269 images (150 ‘control’, and 119 ‘trichostatin A treated’), and were acquired as outlined in (Towers *et al*., 2008). The images of both datasets were JPEG format, and varied in resolution, from 188 x 188 to 1000 x 1000 pixels.

### Clustering analysis

The dimensionality of the raw images was reduced via principal component (PC) analysis (Partridge and Calvo, 1998). The appropriate number of PCs was determined to be 2 by iteratively increasing this number from 1 until we found diminishing returns in the proportion of variance explained (**Fig S1**). *k*-means clustering was performed on the dimensionally-reduced dataset (Ranjan *et al*., 2017), determining the appropriate number for *k* to be 3 by iteratively increasing this number from 1 until we found diminishing returns in the reduction in within-cluster sum of squares (Bholowalia and Kumar, 2014). Haralick image texture features were computed as a feature extraction method (Haralick *et al*., 1973), prior to clustering analysis as above (**Fig S2**).

### Data preprocessing

Preprocessing steps were applied to encourage the trained model to be invariant to image features that are not classifying (e.g. scale, colour) (**Table 1**, **Table S2**, **Fig S3**). Images were converted to grayscale, resizing to 200 x 200 pixels. 200 x 200 is sufficiently small to be easily processed, whilst retaining sufficient spatial resolution to distinguish morphology. Histograms of each image were normalised to brighten images that were too dark and vice versa.

### Data augmentation

The following augmentations were applied in various combinations: Rotation (each image rotated by 36 multiples of 10°), Crop (parameters), Shear (parameters), (Gaussian) blur (parameters), Cutout (DeVries and Taylor, 2017)that aims to reduce the classifier’s reliance on those masked features (DeVries and Taylor, 2017), Möbius transformations (bijective conformal mappings that preserve angles and which may be effective in accounting for user error in microscopy image acquisition, e.g. sample damage during preparation (Zhou *et al*., 2021). Our rotation method enlarged the images on rotation without cutting off any part. This meant that in addition to rotational and colour invariance, scale invariance would be included into the baseline datasets. For the wing classification, we also incorporated flipped images as part of the baseline.

### Traditional classifiers

For each of our Random Forest Classifier (RFC), Support Vector Machine (SVM), and *k*-Nearest Neighbour (KNN) classifiers, we fitted 10 separate models, generating a new training and validation split for each model. Our splitting followed a 80:20 ratio (120 training, 32 validation images).

### Cross-validation

We used a nested cross validation scheme, where at the outset 20% of the dataset was set aside as a stratified test set, i.e. the ratio of labels in the test set reflecting the ratios of the entire datasets (in the brain, a ratio of 0.45:0.54; 10 (early):10 (late) and, in the wing – 0.54:0.44; control:treated). On the remaining data, we employed k-fold cross-validation to compare augmentation/preprocessing performance and avoid overfitting. Briefly, we partitioned the dataset into 10 non-overlapping folds. We then trained the network on folds 2-10 and validated on fold 1. Following this, the network was trained with folds 1 and 3-10, with the second subset used for validation. This proceeded until all folds were used. In this way, we validated the performance of our neural network across the entire dataset. Finally, the DCNNs were evaluated on an independent, (unaugmented) test set, and the highest accuracy DCNNs used for the respective saliency analyses.

### InceptionV3 and ResNet50

For retraining the InceptionV3 (Szegedy *et al*., 2015) and ResNet50 (He *et al*., 2016) networks we used the publicly available ImageNet weights (i.e. the weights of the network which had achieved high performance on ImageNet), then trained between 1 and 500 epochs, halting training if 10 epochs had passed without increasing validation accuracy >0.01%. The number of epochs to pass, and the early stopping threshold, were selected empirically based on the speed at which models that were allowed to train for 500 epochs converged. When this was triggered, we restored the highest scoring weights in training before saving the model. Following (Goodfellow and Bengio, 2016) we inserted a softmax classification layer as the last layer in the model. The softmax activation performs the actual classification by transforming the input between 0 and 1 outputting two values which sum to 1, which effectively define probabilities of the input belonging to each sub-stage. This was necessary as both InceptionV3 and ResNet50 were designed around the ImageNet dataset (1000 classes). We used the optimiser Adam at 10^−5^ (Zhang and Mitliagkas, 2019; Margapuri *et al*., 2020).

### Neural network architecture

The bespoke DNN was based on the Visual Geometry Group (VGG-16) model architecture (Simonyan and Zisserman, 2014) (**Fig 2M**). This architecture involves repeated functional units or VGG ‘blocks’, each comprising a convolutional layer with resolution preservation followed by a max-pooling layer that performs 2x spatial down-sampling. In contrast to VGG-16, we include a single convolutional layer between each max pooling layer. However, we retain the small filter sizes (3 x 3), which help to capture local features, an important step in fine-grained image classification. Between the convolutional and max-pooling layers there is a Rectified Linear Unit (ReLU) activation function (Agarap, 2018). As the actual spatial resolution of the data decreases, the number of filters doubles. Thus, the first layer that receives the 200 x 200 image input has 16 functional units, which is repeated 6 more times resulting in a final convolutional layer with dimensions 4 x 4, with 1024 functional units. This follows a similar pattern to VGG-16, however the largest convolutional layer in VGG-16 is 512 wide, whereas we extend to 1024. Following these blocks, we include 3 fully connected layers (of 1024, 2048, 2048 units), followed by a softmax classification layer; in contrast, VGG-16 uses three wider (4096 units) fully connected layers.

### Training regime

Optimal hyperparameters for our baseline are summarised in **Table S4**. We regularised our network with *L*_2_ regularisation (weight decay), which penalises large weights in a neural network (Goodfellow and Bengio, 2016). The key parameter, *λ*, is a fraction of the sum of the squared weights of the network. As *λ* increases, the loss function value increases. Since a neural network is optimised by minimising the loss function, *L*_2_ regularisation encourages smaller weights and thus less complex models. Our optimal value of 10^−4^ for *λ* was determined by Bayesian optimisation and has been found to be effective in other training image classifiers (Gabas *et al*., 2016). We also used dropout, which randomly turns off neurons in a layer at a given rate (Srivastava, 2013). This discourages individual neurons from becoming dominant, encouraging a classifier with better generalisability. We added a 20% dropout layer between each convolutional and max pooling layer, and a 50% dropout layer before the final classification; these percentages were also determined through Bayesian optimisation. We used the optimiser Adam, with a learning rate of 10^−5^, determined through Bayesian optimisation. We set our range of trialled learning rates to test during optimisation (10^−1^–10^−6^) according to our InceptionV3 / ResNet50 learning rate of 10^−5^.

### Saliency analysis

In the saliency maps, image pixels were generated using the SmoothGrad method and coloured based on whether they contributed more (hot colours) or less (cold colours) towards the output prediction (Smilkov *et al*., 2017). This produced a map of the input features that the network deemed most and least important towards a classification. Saliency maps used test images not involved in model training or validation.

### Software

Image grayscale conversion, resizing, histogram normalisation were implemented using OpenCV (3.4.2.17) and pillow (8.3.1) (Bradski, 2000; Clark, 2015). All augmentations were implemented with imgaug 4.0 and augmentation parameters were randomly selected from ranges given in the provided codes (Jung *et al*., 2020). Clustering was implemented in Python 3.7.12 using scikit-learn 0.24.1 (Pedregosa *et al*., 2011). **Figs S1-2** were generated using seaborn 0.11.2 (Waskom *et al*., 2021) and matplotlib 3.2.2 (Hunter, 2007). Hyperparameter fine-tuning was implemented using keras-tuner 1.1.3 (O’Malley *et al*., 2019). All neural networks were built with Keras 2.10.0 (Chollet and others, 2015) and trained using TensorFlow 2.10.0 (Abadi *et al*., 2015). Neural networks were built and trained using Python 3.6. The models were trained on a mixture of a NVIDIA Tesla V100 GPU, using the HPC system provided by the Joint Academic Data Science Endeavour (JADE) II, and a NVIDIA RTX 4070. Saliency maps were generated using tf-keras-vis 0.8.0 (Yasuhiro, 2021). A full software dependency list is provided at: https://github.com/ianbgroves/chick_embryo_DCNN_classifier.

## Acknowledgments

We thank K. Chinnaiya, E. Manning, and E. Place for providing images of chick embryos, including live-imaging, HCRs and fate-maps used in Figs 1 and 2, and for comments on the manuscript.

## Data availability

All code and datasets from the current study, including a Google Colab notebook to facilitate re-use by users with minimal coding experience, are publicly available from https://github.com/ianbgroves/chick_embryonic_classifier.

## Ethics approval

All experiments used to generate the data were carried out according to the UK Animals (Scientific Procedures) Act 1986. Named Animal Care and Welfare Officers (NACWOs) had oversight of all incubated eggs.

## Authors’ contributions

I.G. Project conception, chick brain data generation, brain data labelling, data analysis, programming, manuscript writing. J.H. Data analysis, programming. D.F. Data analysis, programming. M.T. Chick wing data generation, manuscript review. B.E. Project conception, manuscript review. M.P. Project conception, specification studies, brain data labelling, manuscript writing. A.G.F. Project conception, manuscript writing. All authors read and approved the final manuscript.

## SUPPLEMENTARY INFORMATION

**Fig S1.**
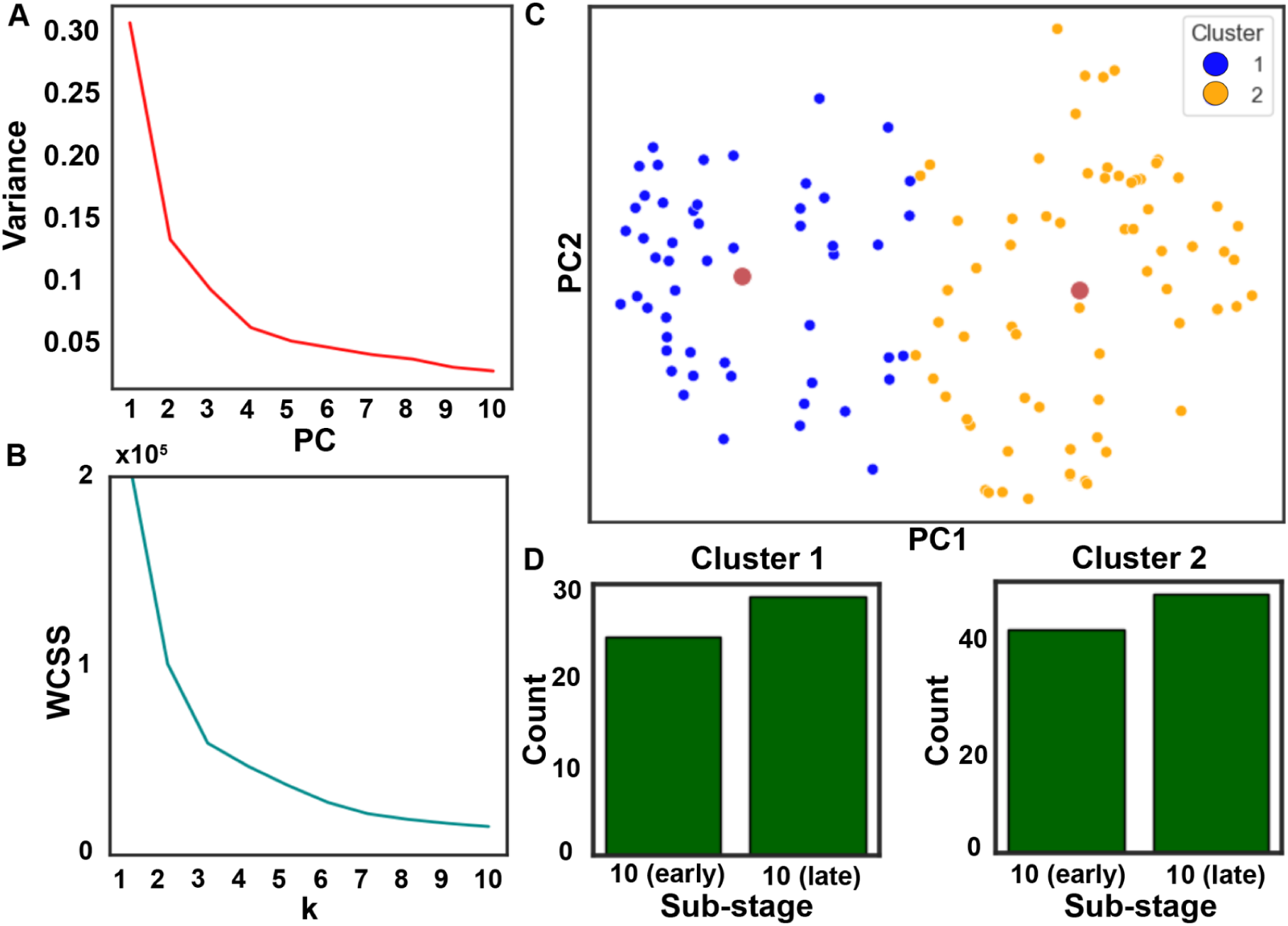
Unsupervised clustering is inaccurate as a biologically relevant classification method. (A) Scree plot for principal component (PC) analysis. The elbow point occurs at 2 PCs, which explain 44% of the variance in the dataset. (B) Plot of the within-cluster sum of squares (WCSS) score from *k*-means clustering with number of *k*. The inflection point occurs at *k*=2. (C) Scatter plot of the 2-means clustered dataset, with centroids (red circles). (D) Number of sub-stages present in each cluster. The number of embryos in each cluster does not match the number of embryos in each labelled sub-stage of the training data. Thus, while this method can sort the data into two populations, these are not developmentally discrete sub-stages.

**Fig S2.**
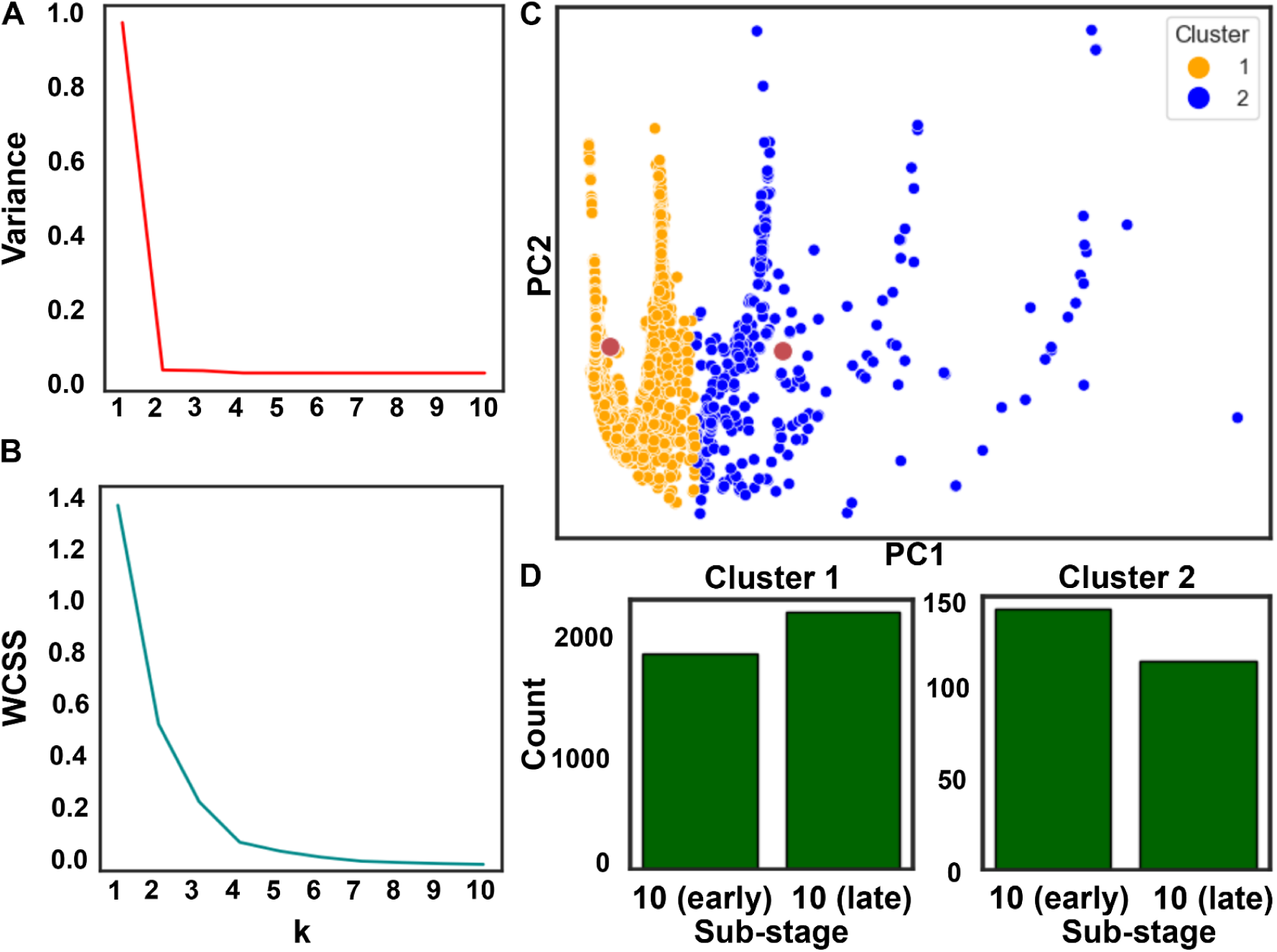
Haralick texture extraction followed by unsupervised clustering is inaccurate as a biologically relevant classification method. (A) Scree plot for principal component analysis (PC) analysis. The elbow point occurs at 2 PCs, which explain 99% of the variance in the dataset. (B) Plot of the within-cluster sum of squares (WCSS) score from *k*-means clustering with number of *k*. The inflection point occurs at *k*=2. (C) Scatter plot of the 2-means clustered haralick features, with centroids (red circles). (D) Number of sub-stages present in each cluster. The number of embryos in each cluster does not match the number of embryos in each labelled sub-stage of the training data. Thus, while this method can sort the data into two populations, these are not developmentally discrete sub-stages.

**Fig S3.**
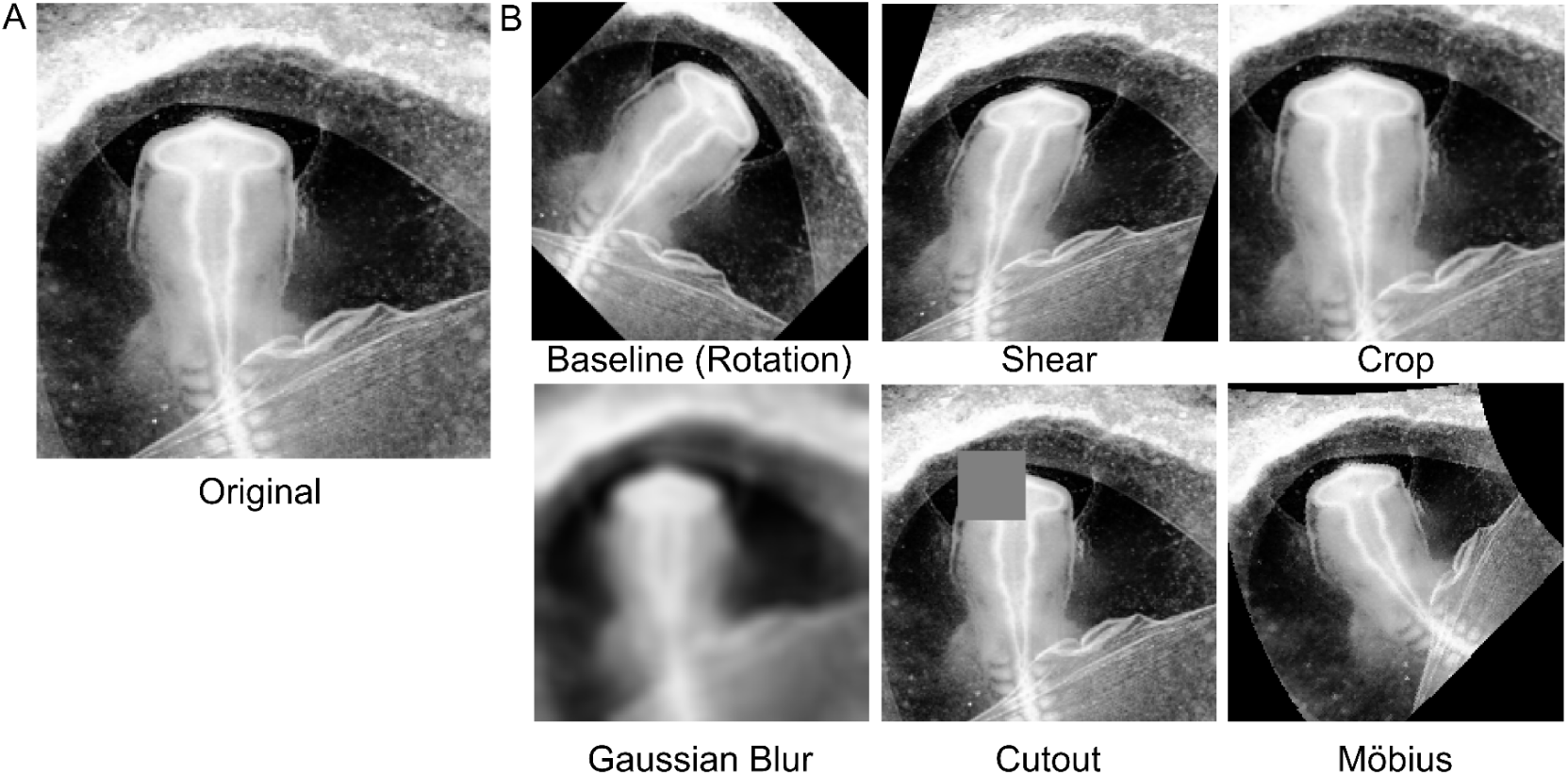
Augmentation regimes. (A) An example image of an embryo from the dataset. (B) the augmentation regimes tested for classifier training involved geometric transformations (e.g. baseline (rotation), shear, crop), photometric (e.g. Gaussian blur), and complex (e.g. cutout, Möbius) augmentations. Augmentations were designed to negate likely artefacts introduced in sample preparation or imaging, such as tissue tearing, which may result in inconsistent variance in e.g. the shapes present in the image. Rotations would negate misaligned samples, shear may negate distortions that could be routinely introduced in sample preparation or imaging. Crop would negate variations in field of view. Gaussian blur may negate variation in focus and/or smooth tissue tearing artefacts. Cutout would reduce over reliance on a single region of the image. Möbius transformations distort shapes but preserve global structure (Zhou *et al.*, 2021), potentially encouraging learning of more general features.

**Table S1.**
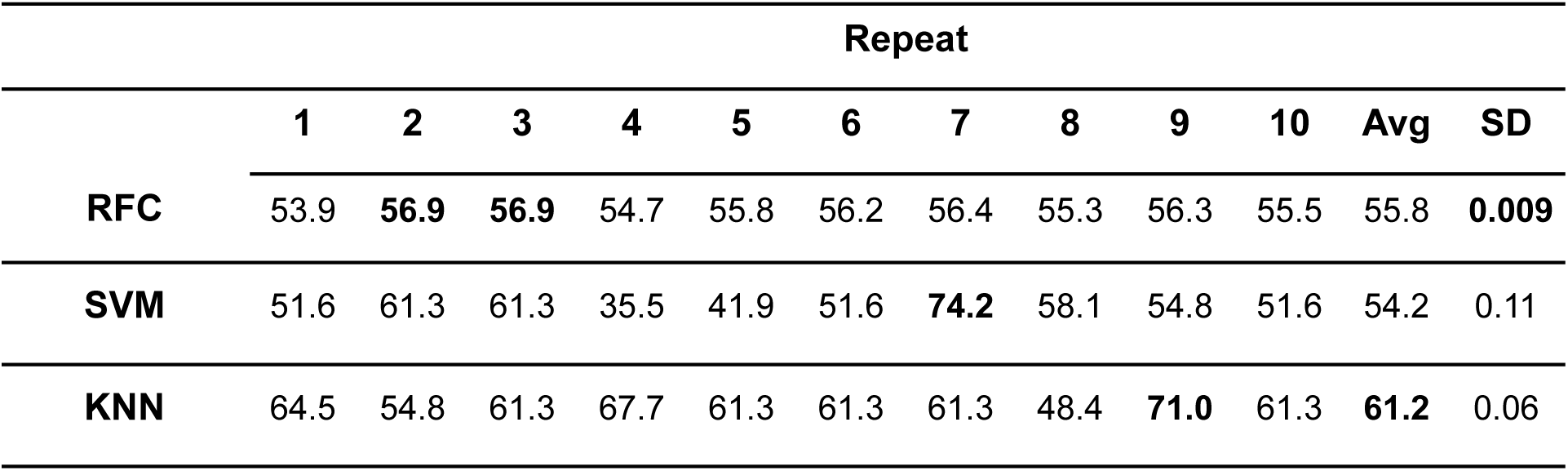
**Traditional machine learning classification on our un-augmented dataset.** Our original dataset was used to fit ten classifiers, and the classification accuracies were determined with a different (80:20) split of training / testing data for each model. RFC; Random forest classifier. SVM; support vector machine, KNN; *k-*nearest neighbours (*k*=3). Highest classification accuracies for each repeat, highest average for each classifier (Avg), and lowest standard deviation (SD) are shown in bold.

**Table S2.**
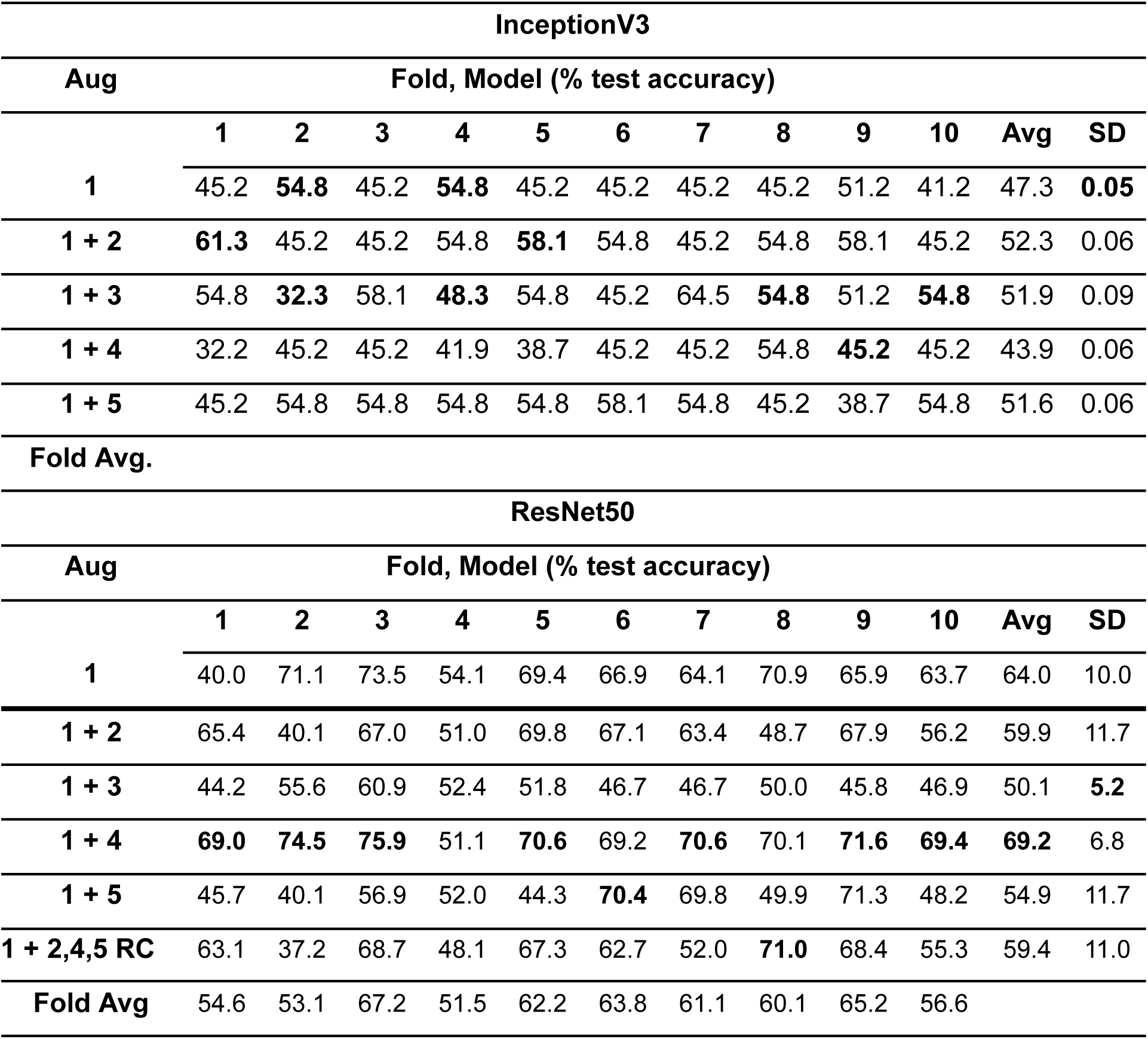
**Augmentation exploration of the dataset using InceptionV3 and ResNet50.** We used *k*-fold cross validation. The individual test accuracies achieved by each network are shown in columns 1-10, and the averages and standard deviation of these accuracies is shown in the rightmost columns. As a baseline processing step, all images were rotated 15 times, at equally spaced degrees. We then tested augmentations on top of this baseline, before a final test in which each image was randomly augmented. Augmentations (Aug) as follows: (1) rotation (baseline); (2) shear; (3) crop; (4) Gaussian blur; (5) cutout; (RC) random combination of rotation + cutout, or shear, or blur. Highest test accuracies for each fold, highest average for each augmentation (Avg), and lowest standard deviation (SD) are shown in bold.

| InceptionV3 |  |  |  |  |  |  |  |  |  |  |  |  |
| --- | --- | --- | --- | --- | --- | --- | --- | --- | --- | --- | --- | --- |
| Aug | Fold, Model (% test accuracy) |  |  |  |  |  |  |  |  |  |  |  |
|  | 1 | 2 | 3 | 4 | 5 | 6 | 7 | 8 | 9 | 10 | Avg | SD |
| **1** | 45.2 | **54.8** | 45.2 | **54.8** | 45.2 | 45.2 | 45.2 | 45.2 | 51.2 | 41.2 | 47.3 | **0.05** |
| **1 + 2** | **61.3** | 45.2 | 45.2 | 54.8 | **58.1** | 54.8 | 45.2 | 54.8 | 58.1 | 45.2 | 52.3 | 0.06 |
| **1 + 3** | 54.8 | **32.3** | 58.1 | **48.3** | 54.8 | 45.2 | 64.5 | **54.8** | 51.2 | **54.8** | 51.9 | 0.09 |
| **1 + 4** | 32.2 | 45.2 | 45.2 | 41.9 | 38.7 | 45.2 | 45.2 | 54.8 | **45.2** | 45.2 | 43.9 | 0.06 |
| **1 + 5** | 45.2 | 54.8 | 54.8 | 54.8 | 54.8 | 58.1 | 54.8 | 45.2 | 38.7 | 54.8 | 51.6 | 0.06 |
| **Fold Avg.** |  |  |  |  |  |  |  |  |  |  |  |  |
| ResNet50 |  |  |  |  |  |  |  |  |  |  |  |  |
| Aug | Fold, Model (% test accuracy) |  |  |  |  |  |  |  |  |  |  |  |
|  | 1 | 2 | 3 | 4 | 5 | 6 | 7 | 8 | 9 | 10 | Avg | SD |
| **1** | 40.0 | 71.1 | 73.5 | 54.1 | 69.4 | 66.9 | 64.1 | 70.9 | 65.9 | 63.7 | 64.0 | 10.0 |
| **1 + 2** | 65.4 | 40.1 | 67.0 | 51.0 | 69.8 | 67.1 | 63.4 | 48.7 | 67.9 | 56.2 | 59.9 | 11.7 |
| **1 + 3** | 44.2 | 55.6 | 60.9 | 52.4 | 51.8 | 46.7 | 46.7 | 50.0 | 45.8 | 46.9 | 50.1 | **5.2** |
| **1 + 4** | **69.0** | **74.5** | **75.9** | 51.1 | **70.6** | 69.2 | **70.6** | 70.1 | **71.6** | **69.4** | **69.2** | 6.8 |
| **1 + 5** | 45.7 | 40.1 | 56.9 | 52.0 | 44.3 | **70.4** | 69.8 | 49.9 | 71.3 | 48.2 | 54.9 | 11.7 |
| **1 + 2,4,5 RC** | 63.1 | 37.2 | 68.7 | 48.1 | 67.3 | 62.7 | 52.0 | **71.0** | 68.4 | 55.3 | 59.4 | 11.0 |
| **Fold Avg** | 54.6 | 53.1 | 67.2 | 51.5 | 62.2 | 63.8 | 61.1 | 60.1 | 65.2 | 56.6 |  |  |

**Table S3.** **Testing of Möbius transformations as data augmentations for the brain dataset.** We used k-fold cross validation. The individual test accuracies achieved by each network are shown in columns 1-10, and the averages (Avg) and standard deviation (SD) of these accuracies is shown in the rightmost columns. As a baseline processing step, all images were rotated 15 times, at equally spaced degrees. 1 + Möbius: the dataset is augmented with our baseline & Möbius transformations. 1 + M:G (10% chance). The dataset is augmented with Gaussian blur with a 10% chance of a Möbius transformation per image.

| Aug | Fold |  |  |  |  |  |  |  |  |  |  |  |
| --- | --- | --- | --- | --- | --- | --- | --- | --- | --- | --- | --- | --- |
|  | 1 | 2 | 3 | 4 | 5 | 6 | 7 | 8 | 9 | 10 | Avg | SD |
| **1 + Möbius** | 55.7 | 57.6 | 47.4 | 49.1 | 49.4 | 41.3 | 38.1 | 57.4 | 43.9 | 66.1 | 50.6 | 8.6 |
| **1 + M:G (10% chance)** | 76.6 | 79.4 | 67.4 | 66.4 | 79.8 | 76.9 | 70.8 | 77.6 | 84.6 | 79.0 | 75.9 | 5.8 |

**Table S4.** **Optimal hyperparameters for our baseline, determined by Bayesian optimisation & empirical selection.** We tested these hyperparameters with the following ranges: Activation function: ReLU-Sigmoid, Batch size: 16-128 – selecting the largest that can fit into the RAM of the GPU, Optimiser: Adam, Adadelta, Adamax, Adagrad, SGD, RMSprop, Layer dropout: 20-30%. Final layer dropout: 50-60%. λ: 10^−3^–10^−4^, Learning rate: 10^−3^–10^−5^, selecting the value/category which was used most by the optimisation algorithm.

| Hyperparameter | Value/category |
| --- | --- |
| Activation function | ReLU |
| Batch size | 32 |
| Layer dropout | 20% |
| Final layer dropout | 50% |
| L<sub>2</sub> regularisation $\lambda$ | $10^{-4}$ |
| Optimiser | Adam |
| Learning rate | $10^{-5}$ |
| Average validation accuracy | 86.2% |
| Min, Max validation accuracy | 78.9%, 96.5% |

## REFERENCES

1. Abadi, M., Agarwal, A., Barham, P., Brevdo, E., Chen, Z., Citro, C., Corrado, G.S., Davis, A., Dean, J., Devin, M., et al. (2015) ‘TensorFlow: Large-Scale Machine Learning on Heterogeneous Systems’. Available at: https://www.tensorflow.org/.

2. Agarap, A.F. (2018) ‘Deep learning using rectified linear units (ReLU)’, arXiv preprint arXiv:1803.08375 [Preprint].

3. Amancio, D.R., Comin, C.H., Casanova, D., Travieso, G., Bruno, O.M., Rodrigues, F.A. and Costa, L. da F. (2014) ‘A systematic comparison of supervised classifiers’, PLOS ONE, 9(4), p. e94137. Available at: https://doi.org/10.1371/journal.pone.0094137.

4. Baltruschat, I.M., Nickisch, H., Grass, M., Knopp, T. and Saalbach, A. (2019) ‘Comparison of deep learning approaches for multi-label chest X-ray classification’, Sci. Rep., 9(1), pp. 1–10.

5. Bergstra, J. and Bengio, Y. (2012) ‘Random search for hyper-parameter optimization.’, J. Mach. Learn. Res., 13(2).

6. Bholowalia, P. and Kumar, A. (2014) ‘EBK-means: A clustering technique based on elbow method and k-means in WSN’, Int. J. Comput. Appl., 105(9).

7. Boehm, B., Rautschka, M., Quintana, L., Raspopovic, J., Jan, Ž. and Sharpe, J. (2011) ‘A landmark-free morphometric staging system for the mouse limb bud’, Development, 138, pp. 1227–1234. Available at: https://doi.org/10.1242/dev.057547.

8. Bouyahya, A., El Omari, N., Bakha, M., Aanniz, T., El Menyiy, N., El Hachlafi, N., El Baaboua, A., El-Shazly, M., Alshahrani, M.M., Al Awadh, A.A., et al. (2022) ‘Pharmacological properties of trichostatin A, focusing on the anticancer potential: a comprehensive review’, Pharmaceuticals, 15(10), p. 1235. Available at: https://doi.org/10.3390/ph15101235.

9. Bradski, G. (2000) ‘The OpenCV Library’, Dr. Dobb’s Journal of Software Tools [Preprint].

10. Chinnaiya, K., Burbridge, S., Jones, A., Kim, D.W., Place, E., Manning, E., Groves, I., Sun, C., Towers, M., Blackshaw, S., et al. (2023) ‘A neuroepithelial wave of BMP signalling drives anteroposterior specification of the tuberal hypothalamus’, eLife. Edited by E. Knust and M.E. Bronner, 12, p. e83133. Available at: https://doi.org/10.7554/eLife.83133.

11. Chollet, F. and others (2015) ‘Keras’. Available at: https://keras.io.

12. Clark, A. (2015) ‘Pillow (PIL Fork) Documentation’. readthedocs. Available at: https://buildmedia.readthedocs.org/media/pdf/pillow/latest/pillow.pdf.

13. Cutrale, F., Fraser, S.E. and Trinh, L.A. (2019) ‘Imaging, visualization, and computation in developmental biology’, Annu. Rev. Biomed. Data Sci., 2, pp. 223–251.

14. Deng, J., Dong, W., Socher, R., Li, L.-J., Li, K. and Fei-Fei, L. (2009) ‘ImageNet: A large-scale hierarchical image database’, in IEEE Conference on Computer Vision and Pattern Recognition. IEEE, pp. 248–255.

15. DeVries, T. and Taylor, G.W. (2017) ‘Improved regularization of convolutional neural networks with cutout’, arXiv preprint arXiv:1708.*04552*, pp. 1–8.

16. Ding, C. and He, X. (2004) ‘K-means clustering via principal component analysis’, in Proceedings of the twenty-first international conference on Machine learning. New York, NY, USA: Association for Computing Machinery (ICML ’04), p. 29. Available at: https://doi.org/10.1145/1015330.1015408.

17. Fu, T., Towers, M. and Placzek, M.A. (2017) ‘Fgf10+ progenitors give rise to the chick hypothalamus by rostral and caudal growth and differentiation’, Development, 144, pp. 3278–88. Available at: https://doi.org/10.1242/dev.153379.

18. Gabas, A., Corona, E., Alenyà, G. and Torras, C. (2016) ‘Robot-aided cloth classification using depth information and CNNs’, in International Conference on Articulated Motion and Deformable Objects. Springer, pp. 16–23.

19. Goodfellow, I. and Bengio, A., and Y. Courville (2016) Deep Learning. MIT Press.

61. Hamburger, V. and Hamilton, H.L. (1951) ‘A series of normal stages in the development of the chick embryo’, J. Morphol., 88, pp. 49–92. Available at: https://doi.org/10.1002/aja.1001950404.

20. Haralick, R., Shanmugam, K. and Dinstein, I. (1973) ‘Textural features for image classification’, IEEE Trans. Syst. Man Cybern. [Preprint]. Available at: https://doi.org/10.1109/TSMC.1973.4309314.

21. He, K., Zhang, X., Ren, S. and Sun, J. (2016) ‘Deep residual learning for image recognition’, in Proceedings of IEEE Computer Society Conference on Computer Vision and Pattern Recognition, pp. 770–778.

22. Hunter, J.D. (2007) ‘Matplotlib: A 2D graphics environment’, Comput. Sci. Eng., 9, pp. 90–95. Available at: https://doi.org/10.1109/MCSE.2007.55.

23. Ishaq, O., Sadanandan, S.K. and Wählby, C. (2017) ‘Deep Fish’, SLAS Discovery, 22(1), pp. 102–107. Available at: https://doi.org/10.1177/1087057116667894.

24. Jacquemet, G. (2021) ‘Deep learning to analyse microscopy images’, Biochem., 43, pp. 60–64. Available at: https://doi.org/10.1042/bio_2021_167.

25. Jung, A.B., Wada, K., Crall, J., Tanaka, S., Graving, J., Reinders, C., Yadav, S., Banerjee, J., Vecsei, G., Kraft, A., et al. (2020) ‘imgaug’. Available at: https://github.com/aleju/imgaug.

26. Kim, D.W., Place, E., Chinnaiya, K., Manning, E., Sun, C., Dai, W., Groves, I., Ohyama, K., Burbridge, S., Placzek, M., et al. (2022) ‘Single-cell analysis of early chick hypothalamic development reveals that hypothalamic cells are induced from prethalamic-like progenitors’, Cell Rep., 38(3), p. 110251. Available at: https://doi.org/10.1016/j.celrep.2021.110251.

27. Kora, P., Ooi, C.P., Faust, O., Raghavendra, U., Gudigar, A., Chan, W.Y., Meenakshi, K., Swaraja, K., Plawiak, P. and Rajendra Acharya, U. (2022) ‘Transfer learning techniques for medical image analysis: A review’, Biocybern. Biomed. Eng., 42(1), pp. 79–107. Available at: https://doi.org/10.1016/j.bbe.2021.11.004.

28. LeCun, Y., Bengio, Y. and Hinton, G. (2015) ‘Deep learning’, Nature, 521(7553), pp. 436–444. Available at: https://doi.org/10.1038/nature14539.

29. LeCun, Y., Bottou, L., Bengio, Y. and Haffner, P. (1998) ‘Gradient-based learning applied to document recognition’, Proc. IEEE, 86(11), pp. 2278–2324. Available at: https://doi.org/10.1109/5.726791.

30. Margapuri, V., Lavezzi, G., Stewart, R. and Wagner, D. (2020) ‘Bombus Species Image Classification’, *arXiv preprint arXiv:2006.11374* [Preprint].

31. Musy, M., Flaherty, K., Raspopovic, A., J. and Robert-Moreno, Richtsmeier, J.T. and Sharpe, J. (2018) ‘A quantitative method for staging mouse embryos based on limb morphometry’, Development, 145, p. dev154856. Available at: https://doi.org/10.1242/dev.154856.

32. Newgreen, D.F. and Erickson, C.A. (1986) ‘The migration of neural crest cells’, Int. Rev. Cyt., 103, pp. 89–145. Available at: https://doi.org/10.1016/S0074-7696(08)60834-7.

33. Ohyama, K., Ellis, P., Kimura, S. and Placzek, M. (2005) ‘Directed differentiation of neural cells to hypothalamic dopaminergic neurons’, Development, 132(23), pp. 5185–5197. Available at: https://doi.org/10.1242/dev.02094.

34. O’Malley, T., Bursztein, E., Long, J., Chollet, F., Jin, H., Invernizzi, L., and others (2019) ‘Keras Tuner’. Available at: https://github.com/keras-team/keras-tuner.

35. O’Rahilly, R. and Müller, F. (2010) ‘Developmental stages in human embryos: revised and new measurements’, Cells Tissues Organs, 192(2), pp. 73–84. Available at: https://doi.org/10.1159/000289817.

36. Palmeirim, I., Henrique, D., Ish-Horowicz, D. and Pourquié, O. (1997) ‘Avian hairy gene expression identifies a molecular clock linked to vertebrate segmentation and somitogenesis’, Cell, 91, pp. 639–648. Available at: https://doi.org/10.1016/S0092-8674(00)80451-1.

37. Panwar, H., Gupta, P.K., Siddiqui, M.K., Morales-Menendez, R., Bhardwaj, P. and Singh, V. (2020) ‘A deep learning and grad-CAM based color visualization approach for fast detection of COVID-19 cases using chest X-ray and CT-Scan images’, Chaos Solitons Fractals, 140, p. 110190. Available at: https://doi.org/10.1016/j.chaos.2020.110190.

38. Partridge, M. and Calvo, R.A. (1998) ‘Fast dimensionality reduction and simple PCA’, Intell. Data Anal., 2(3), pp. 203–214. Available at: https://doi.org/10.1016/S1088-467X(98)00024-9.

39. Pedregosa, F., Varoquaux, G., Gramfort, A., Michel, V., Thirion, B., Grisel, O., Blondel, M., Prettenhofer, P., Weiss, R., Dubourg, V., et al. (2011) ‘Scikit-learn: machine learning in Python’, J. Mach. Learn. Res., 12, pp. 2825–2830.

40. Pond, A.J.R., Hwang, S., Verd, B. and Steventon, B. (2021) ‘A deep learning approach for staging embryonic tissue isolates with small data’, PLOS ONE, 16, p. e0244151. Available at: https://doi.org/10.1371/journal.pone.0244151.

41. Ranjan, S., Nayak, D.R., Kumar, K.S., Dash, R. and Majhi, B. (2017) ‘Hyperspectral image classification: A k-means clustering based approach’, in 4th International Conference on Advanced Computing and Communication Systems, pp. 1–7. Available at: https://doi.org/10.1109/ICACCS.2017.8014707.

42. Rosin, P.L. and Fierens, F. (1995) ‘Improving neural network generalisation’, in International Geoscience and Remote Sensing Symposium. IEEE, pp. 1255–1257.

43. Sáenz-Ponce, N., Mitgutsch, C. and del Pino, E.M. (2012) ‘Variation in the schedules of somite and neural development in frogs’, Proc. Natl. Acad. Sci. U.S.A., 109, pp. 20503–20507. Available at: https://doi.org/10.1073/pnas.1219307110.

44. Shin, S., Lee, Y., Kim, M., Park, J., Lee, S. and Min, K. (2020) ‘Deep neural network model with Bayesian hyperparameter optimization for prediction of NOx at transient conditions in a diesel engine’, Eng. Appl. Artif. Intell., 94, p. 103761. Available at: https://doi.org/10.1016/j.engappai.2020.103761.

45. Simard, P.Y., Steinkraus, D. and Platt, J.C. (2003) ‘Best practices for convolutional neural networks applied to visual document analysis’, in Icdar.

46. Simonyan, K., Vedaldi, A. and Zisserman, A. (2014) ‘Deep inside convolutional networks: visualising image classification models and saliency maps’, arXiv:1312.6034 [cs] [Preprint]. Available at: http://arxiv.org/abs/1312.6034 (Accessed: 17 February 2022).

47. Simonyan, K. and Zisserman, A. (2014) ‘Very deep convolutional networks for large-scale image recognition’, arXiv preprint arXiv:1409.1556 [Preprint].

48. Smilkov, D., Thorat, N., Kim, B., Viégas, F. and Wattenberg, M. (2017) ‘Smoothgrad: removing noise by adding noise’, *arXiv preprint arXiv:1706.03825* [Preprint].

49. Srivastava, N. (2013) ‘Improving neural networks with dropout’, University of Toronto, 182(566), p. 7.

50. Stern, C.D. (2018) ‘Staging tables for avian embryos: a little history’, Int. J. Dev. Biol., 62, pp. 43–48. Available at: https://doi.org/10.1387/ijdb.170299cs.

51. Szegedy, C., Liu, W., Jia, Y., Sermanet, P., Reed, S., Anguelov, D., Erhan, D., Vanhoucke, V. and Rabinovich, A. (2015) ‘Going deeper with convolutions’, in Proceedings of IEEE Computer Society Conference on Computer Vision and Pattern Recognition, pp. 1–9.

52. Theiler, K. (2013) *The House Mouse: Atlas of Embryonic Development*. Springer Science & Business Media.

53. Thompson, N.C., Greenewald, K., Lee, K. and Manso, G.F. (2020) ‘The computational limits of deep learning’, *arXiv preprint arXiv:2007.05558* [Preprint].

54. Towers, M., Mahood, R., Yin, Y. and Tickle, C. (2008) ‘Integration of growth and specification in chick wing digit-patterning’, Nature, 452(7189), pp. 882–886. Available at: https://doi.org/10.1038/nature06718.

55. Waskom, M., Gelbart, M., Botvinnik, O., Ostblom, J., Hobson, P., Lukauskas, S., Gemperline, D.C., Augspurger, T., Halchenko, Y., Warmenhoven, J., et al. (2021) ‘mwaskom/seaborn: v0.11.2 (August 2021)’. Zenodo. Available at: https://doi.org/10.5281/zenodo.5205191.

56. Yamashita, R., Nishio, M., Do, R.K.G. and Togashi, K. (2018) ‘Convolutional neural networks: an overview and application in radiology’, Insights into Imaging, 9(4), pp. 611–629. Available at: https://doi.org/10.1007/s13244-018-0639-9.

57. Yasuhiro, K. (2021) ‘tf-keras-vis’, *GitHub repository*. GitHub. Available at: https://github.com/keisen/tf-keras-vis.

58. Zeiler, M.D. and Fergus, R. (2014) ‘Visualizing and understanding convolutional networks’, in D. Fleet, T. Pajdla, B. Schiele, and T. Tuytelaars (eds) Computer Vision – ECCV 2014. Cham: Springer International Publishing (Lecture Notes in Computer Science), pp. 818–833. Available at: https://doi.org/10.1007/978-3-319-10590-1_53.

59. Zhang, J. and Mitliagkas, I. (2019) ‘YellowFin and the art of momentum tuning’, Proceedings of Machine Learning and Systems, 1, pp. 289–308.

60. Zhou, S., Zhang, J., Jiang, H., Lundh, T. and Ng, A.Y. (2021) ‘Data augmentation with Mobius transformations’, Mach. Learn.: Sci. Technol., 2, p. 025016. Available at: https://doi.org/10.1088/2632-2153/abd615.

